## Supporting Information for "DNA i-motif levels are overwhelmingly depleted in living human cells: insights from in-cell NMR"

**Table S1**: List of DNA constructs used in this study.

| Name | Sequence (5' → 3') |
|---|---|
| hybrid-ds/hT121-6 FW | GCT TCT AGT CAA TCC CCC CTC CCC CCT TCC CCC CTC CCC CC |
| FAM_ hybrid-ds/hT121-6 FW | [6FAM] GCT TCT AGT CAA TCC CCC CTC CCC CCT TCC CCC CTC CCC CC |
| hybrid-ds/hTel FW | GCT TCT AGT CAA T CCC TAA CCC TAA CCC TAA CCC |
| FAM_ hybrid-ds/hTel FW | [6FAM] GCT TCT AGT CAA T CCC TAA CCC TAA CCC TAA CCC |
| hybrid-ds/hPDGFa FW | GCT TCT AGT CAA TCC GCG CCC CTC CCC CGC CCC CGC CCC CGC CCC CCC CCC CCC |
| FAM_ hybrid-ds/hPDGFa FW | [6FAM] GCT TCT AGT CAA TCC GCG CCC CTC CCC CGC CCC CGC CCC CGC CCC CCC CCC CCC |
| hybrid-ds/hRAD17 FW | GCT TCT AGT CAA TCC ACC CCC CCC CGC CCC CCC CCG GA |
| FAM_hybrid-ds/hRAD17 FW | [6FAM] GCT TCT AGT CAA TCC ACC CCC CCC CGC CCC CCC CCG GA |
| hybrid-ds RV | TTG ACT AGA AGC |
| hybrid-ds FW | GCT TCT AGT CAA |
| hBcl-2 | CAG CCC CGC TCC CGC CCC CTT CCT CCC GCG CCC GCC CCT |
| FAM_hBcl-2 | [6FAM] CAG CCC CGC TCC CGC CCC CTT CCT CCC GCG CCC GCC CCT |
| Cy3_hT121-6 | [Cyanine3] CCC CCC TCC CCC CTT CCC CCC TCC CCC C |
| Cy3_hTel | [Cyanine3] CCC TAA CCC TAA CCC TAA CCC |
| Cy3_hPDGFa | [Cyanine3] CCG CGC CCC TCC CCC GCC CCC GCC CCC GCC CCC CCC CCC CC |
| Cy3_hRAD17 | [Cyanine3] CCA CCC CCC CCC GCC CCC CCC CGG A |
| Cy3_hBcl-2 | [Cyanine3] CAG CCC CGC TCC CGC CCC CTT CCT CCC GCG CCC GCC CCT |
| Cy3_cMycG | [Cyanine3] TGG GGA GGG TGG GGA GGG TGG GGA AGG TGG GGA GAA GA |
| Cy3_hTelG | [Cyanine3] GGG TTA GGG TTA GGG TTA GGG |

### Panel I

| | iM / **hybrid-ds/iM** | | | | |
|---|---|---|---|---|---|
| | | | Tm (°C) | | |
| iM | **pH 7** | **pH 6.5** | **pH 6** | **pH 5.5** | **pH 5** |
| hTel | n.d. / **n.d.** | 15.9 ± 0.5 / **13.9 ± 0.9** | 26.5 ± 0.3 / **29.2 ± 0.3** | 40.5 ± 0.5 / **42.2 ± 0.4** | 51.3 ± 0.2 / **53.4 ± 0.2** |
| hT121-6 | 24.9 ± 0.5 / **28.0 ± 0.4** | 36.6 ± 0.5 **/28.0 ± 0.4** | 48.3 ± 0.2 / **28.0 ± 0.4** | 63.6 ± 0.3 / **28.0 ± 0.4** | 74.3 ± 0.7 / **28.0 ± 0.4** |
| hPDGFa | 28.8 ± 1.2 / **27.6 ± 0.9** | 37.7 ± 0.3 / **37.4 ± 0.3** | 50.8 ± 0.5 / **49.8 ± 0.7** | 65.7 ± 1.2 / **64.7 ± 0.8** | - |
| hRAD17 | 33.8 ± 0.5 / **34.9 ± 0.6** | 45.1 ± 0.7 / **44.3 ± 1.2** | 55.3 ± 1.0 / **54.2 ± 2.5** | - | - |
| hBcl-2 | 12.9 ± 0.5 | 26.5 ± 0.2 | 38.1 ± 0.2 | 52.2 ± 0.4 | - |

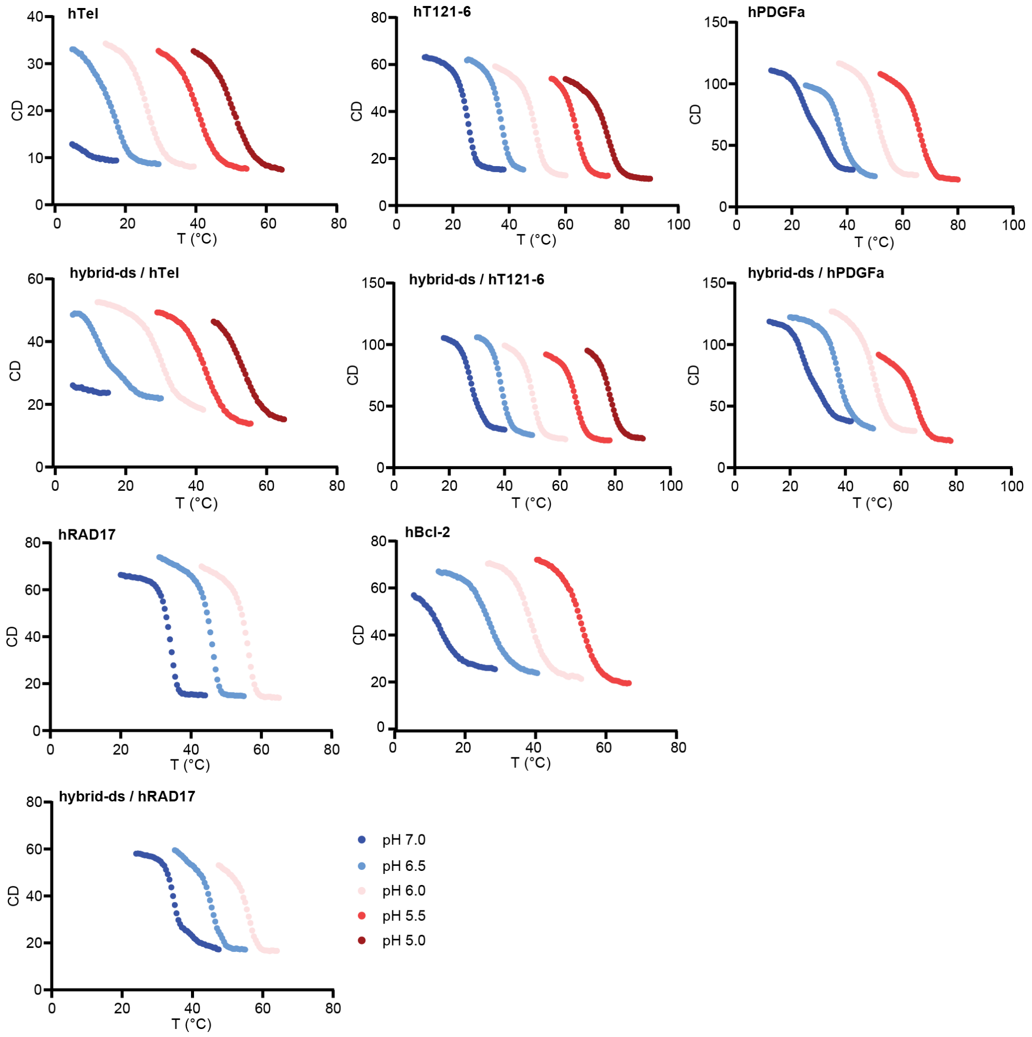

#### Panel II

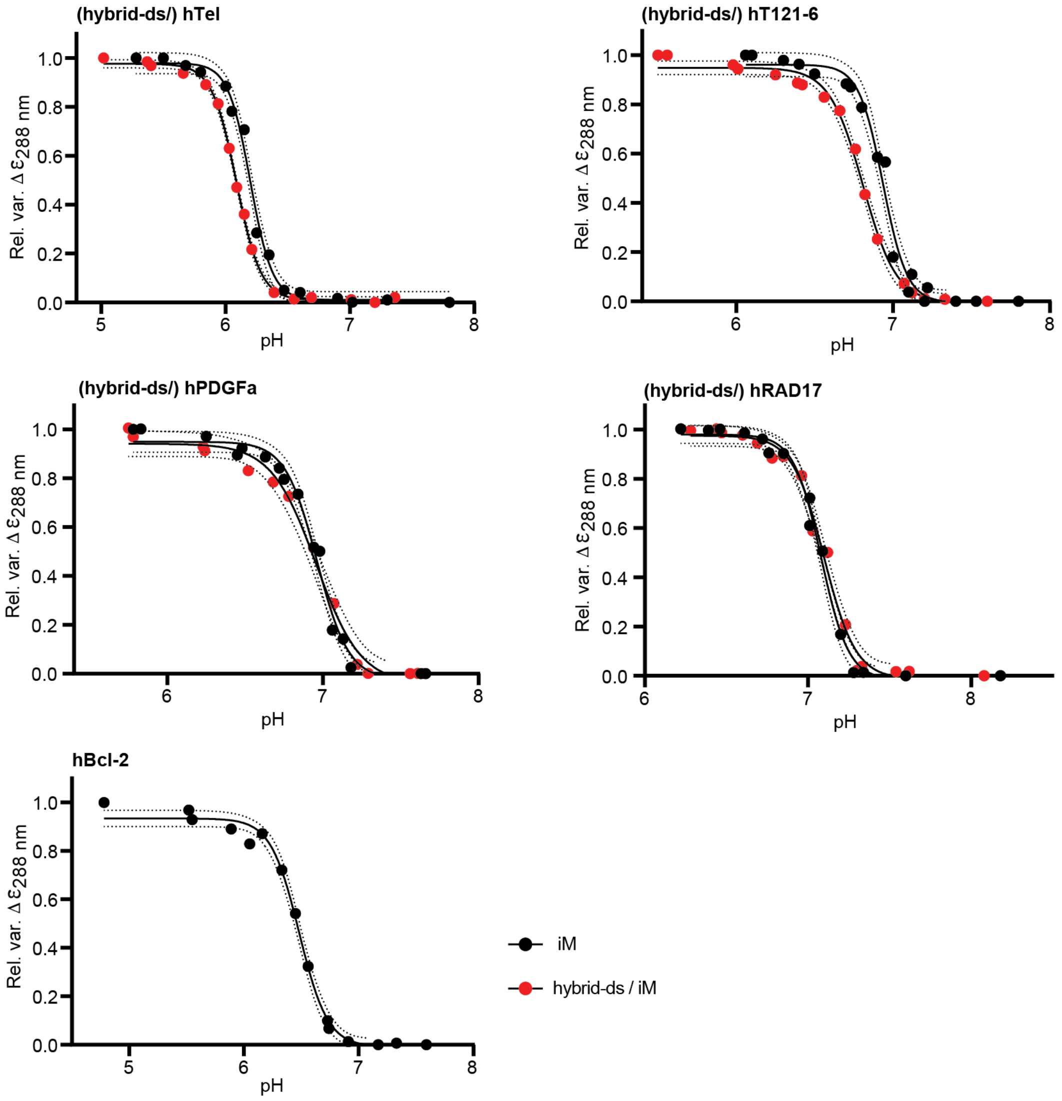


**Fig. S1. Panel I)** Assessment of the thermal stability of iM and hybrid-ds/iM constructs at different pH *in vitro*. The $T_m^{in-vitro}$ values were derived from CD melting profiles of iM and hybrid-ds/iM constructs measured at different pH (as indicated). Each sample was prepared in IC-cacodylate-based buffer (25 mM sodium cacodylate, 10.5 mM NaCl, 110 mM KCl, 1 mM $MgCl_2$, 130 nM $CaCl_2$). The reported error corresponds to the standard error of the mean for each triplicate experiment. **Panel II**) Assessment of pH-dependency of t iM and hybrid-ds/iM constructs. The graphs show the CD signal variations (monitored at 288 nm) as a function of the pH change. Black and red dots correspond to the pH titration for the iM and hybrid-ds/iM, respectively. The measurements were performed at 20 °C in IC buffer (25 mM potassium phosphate, 10.5 mM NaCl, 110 mM KCl, 1 mM $MgCl_2$, 130 nM $CaCl_2$). The panels represent the fitted curves as solid lines, while dotted lines indicate the 95% confidence interval of the fit. The derived $pH_T^{in-vitro}$ values are reported in Table 1.

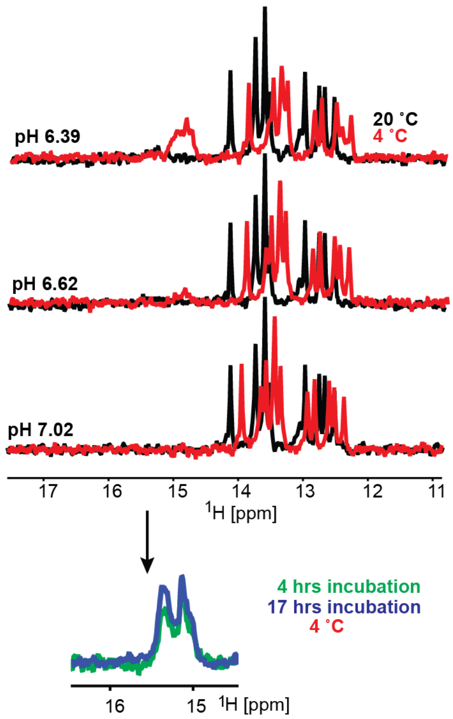


**Fig. S2.** Overlays of the imino regions of 1D $^1$H NMR spectra of hybrid-ds/hTel acquired *in vitro* (IC buffer: 25 mM $KPO_i$, 10.5 mM NaCl, 110 mM KCl, 1 mM $MgCl_2$, 130 nM $CaCl_2$) at 4 °C (red) and 20 °C (black), as a function of pH and the time at 4 °C (green and blue). Note: The temperature-induced chemical shift changes observed for Watson-Crick signals (12-14,5 ppm) are results of (de)stabilization of the double-helical region by temperature alteration (dynamical averaging).

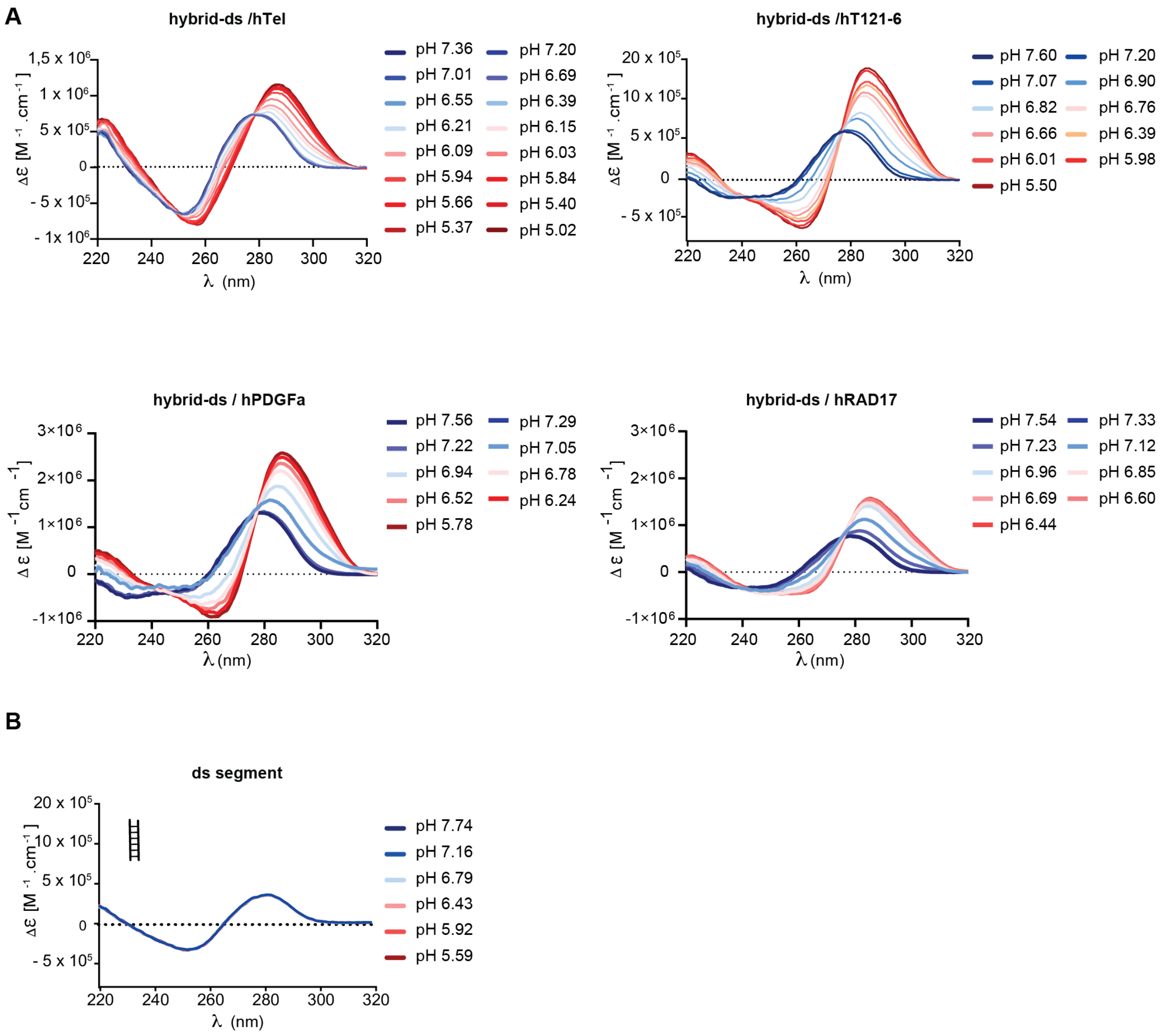


**Fig. S3**. CD spectra of hybrid-ds/iM (A) and isolated ds segment (B) acquired as a function of pH at room temperature *in vitro* (IC buffer: 25 mM $KPO_i$, 10.5 mM NaCl, 110 mM KCl, 1 mM $MgCl_2$, 130 nM $CaCl_2$).

#### Panel I

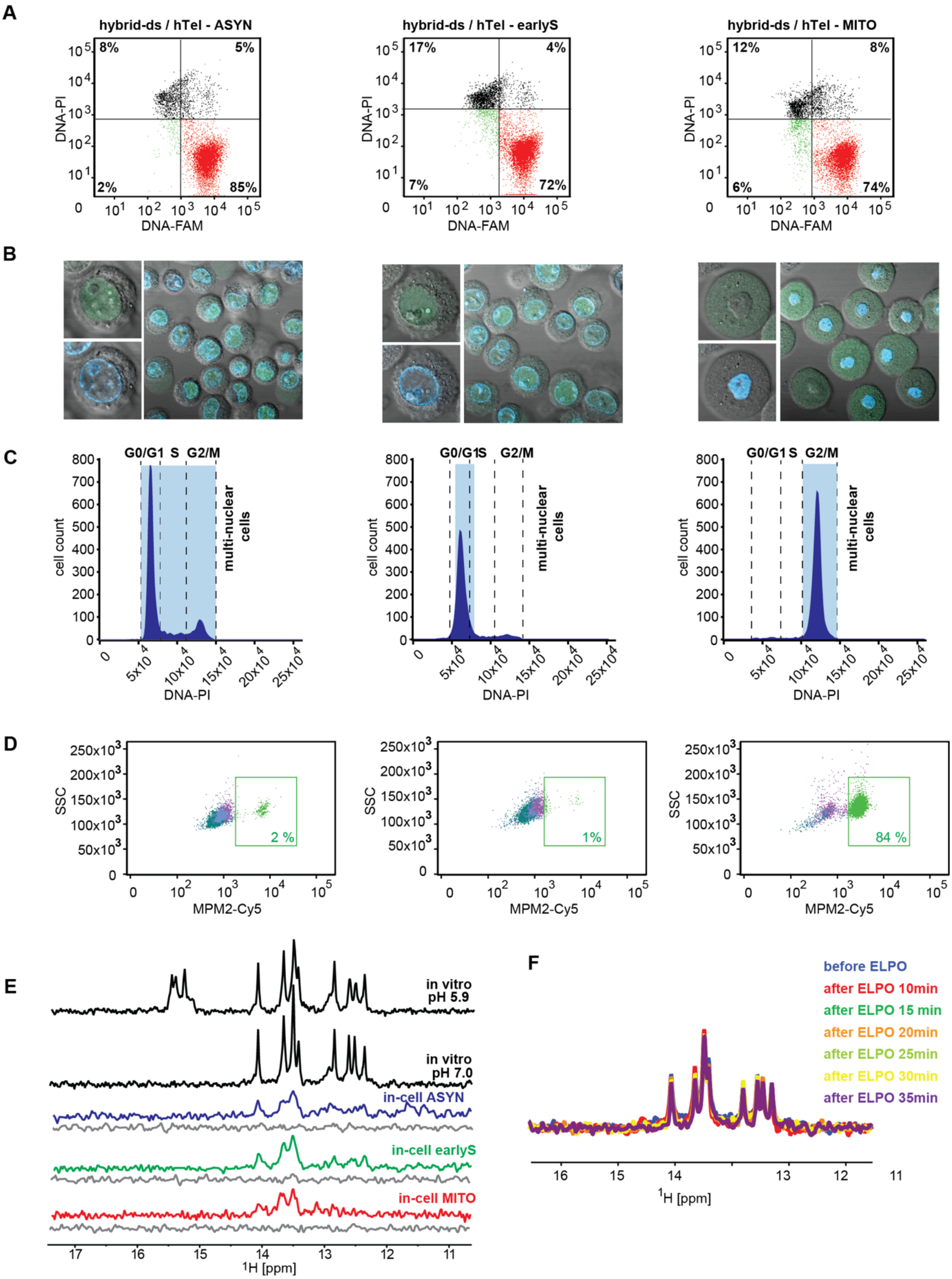

A
hybrid-ds / hTel - ASYN
hybrid-ds / hTel - earlyS
hybrid-ds / hTel - MITO
DNA-PI
DNA-FAM
8%
5%
2%
85%
17%
4%
7%
72%
12%
8%
6%
74%
B
C
G0/G1
S
G2/M
multi-nuclear cells
cell count
DNA-PI
D
SSC
MPM2-Cy5
2 %
1%
84 %
E
in vitro pH 5.9
in vitro pH 7.0
in-cell ASYN
in-cell earlyS
in-cell MITO
$^1$H [ppm]
F
before ELPO
after ELPO 10min
after ELPO 15 min
after ELPO 20min
after ELPO 25min
after ELPO 30min
after ELPO 35min
$^1$H [ppm]

#### Panel II

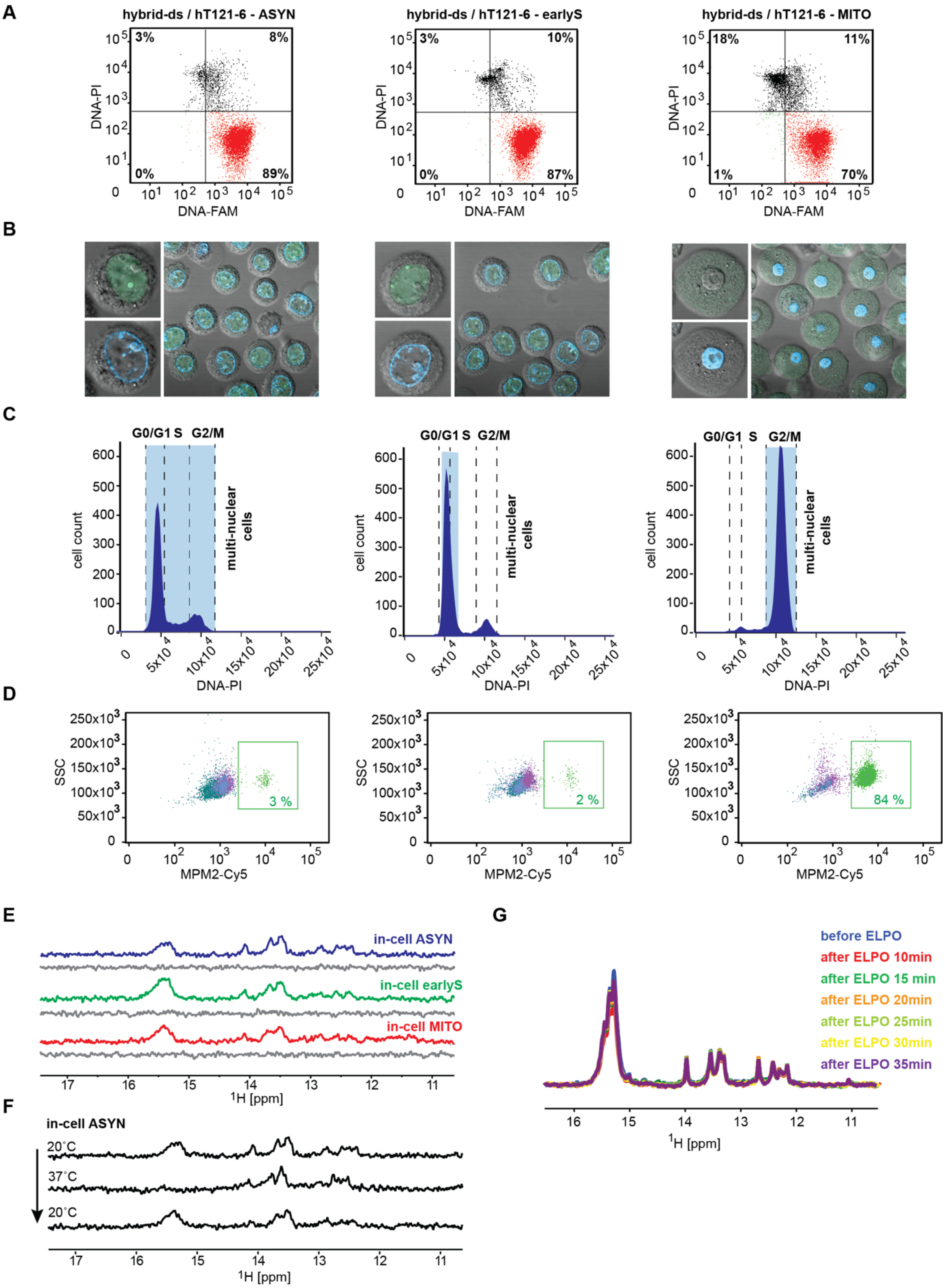
A
hybrid-ds / hT121-6 - ASYN
hybrid-ds / hT121-6 - earlyS
hybrid-ds / hT121-6 - MITO
3%
8%
0%
89%
3%
10%
0%
87%
18%
11%
1%
70%
DNA-PI
DNA-FAM
B
C
G0/G1 S G2/M
multi-nuclear cells
cell count
DNA-PI
D
SSC
MPM2-Cy5
3 %
2 %
84 %
E
in-cell ASYN
in-cell earlyS
in-cell MITO
1H [ppm]
F
in-cell ASYN
20°C
37°C
20°C
1H [ppm]
G
before ELPO
after ELPO 10min
after ELPO 15 min
after ELPO 20min
after ELPO 25min
after ELPO 30min
after ELPO 35min
1H [ppm]

**Fig. S4**. (**A**) Double-staining (DNA-PI/DNA-FAM) FCM analysis (post-in-cell NMR acquisition) of asynchronous HeLa cells (left), HeLa cells synchronized in early-S (middle) and M- (right) cell-cycle phases transfected with (FAM-)hybrid-ds/hTel (**Panel I**) and (FAM-)hybrid-ds/hT121-6 (**Panell II**). The FCM plots indicate the percentages of viable nontransfected cells, viable hybrid-ds/iM DNA-containing cells, dead/compromised nontransfected cells, and dead/compromised cells transfected with hybrid-ds/iMs in the bottom-left, bottom-right, top-left, and top-right quadrants, respectively. (**B**) and (**C**) show confocal microscopy images and propidium iodide (PI) DNA content staining of asynchronous HeLa cells (left), HeLa cells synchronized in early-S (middle), and M- (right) cells transfected with (FAM-)hybrid-ds/hTel (**Panel I**) and (FAM-)hybrid-ds/hT121-6 (**Panell II**), respectively. Green and blue colors in (**B**) indicate the localization of the introduced hybrid-ds/iM and cell nucleus, respectively. (**D**) MPM2-Cy5 staining of asynchronous HeLa cells (left), HeLa cells synchronized in early-S (middle) and M- (right) cells transfected with (FAM-)hybrid-ds/hTel (**Panel I**) and (FAM-)hybrid-ds/hT121-6 (**Panell II**), respectively. (**E**) shows imino regions of 1D $^1$H NMR spectra of hybrid-ds/hTel (**Panel I**) and hybrid-ds/hT121-6 (**Panel II**) acquired in IC buffer (black), and in asyn- (blue), early-S (green) and M-(red) synchronized cells at 20 °C. The corresponding region of the 1D $^1$H NMR spectra of the extracellular fluid taken from the samples after in-cell NMR spectra acquisition is shown in gray (leakage control). (**F – Panel I** and **G - Panel II**) show an overlay of imino regions of 1D $^1$H NMR spectra of hybrid-ds/hTel and hybrid-ds/hT121-6 before and after applying the ”mock” electroporation pulse. (**F – Panel II**) shows imino regions of 1D $^1$H NMR spectra of hybrid-ds/hT121-6 acquired as a function of the temperature (indicated) in suspension of asynchronous HeLa cells.

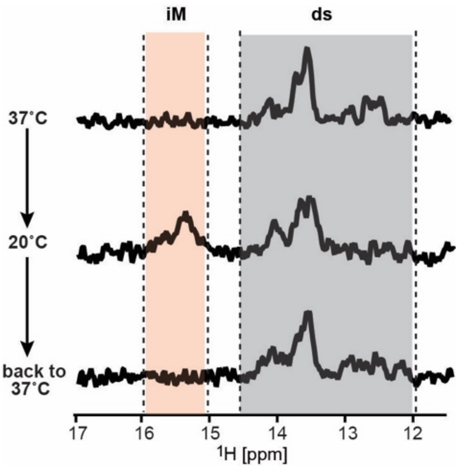


**Fig. S5**. Imino regions of the 1D $^1$H NMR spectra of hybrid-ds/hT121-6 acquired as a function of the temperature (indicated) in low-melting agarose immobilized asynchronous HeLa cells under conditions of the constant media flow (bioreactor) with the acquisition time window of 30 min.

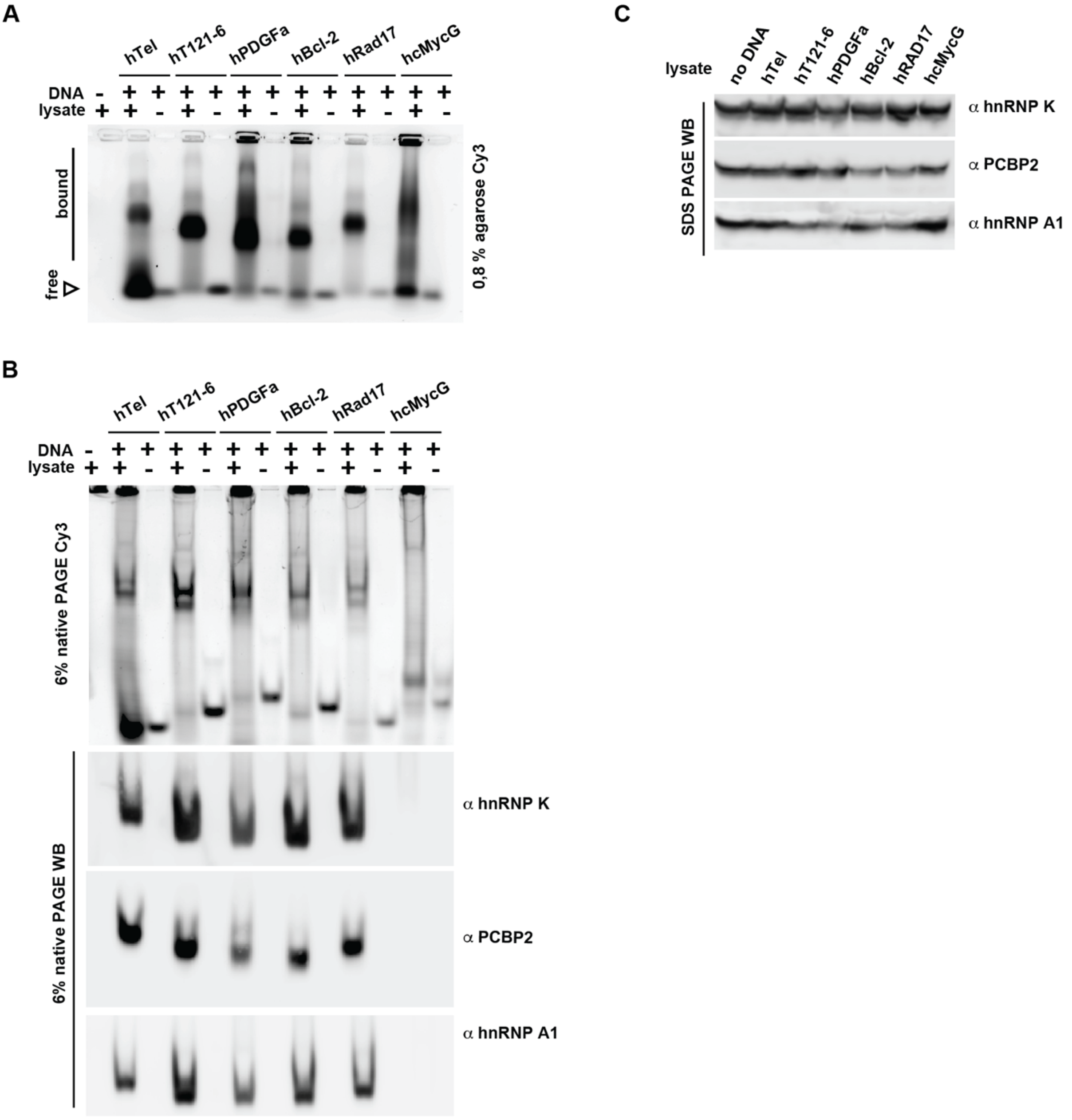


**Fig. S6.** Protein binding to DNA oligonucleotides. (**A**) Electrophoretic migration of the indicated Cy3-labelled oligonucleotides in agarose gel, in the absence and the presence of lysates from HeLa cells, visualized via the Cy3-fluorescence. (**B**) Native PAGE of the indicated Cy3-labelled oligonucleotides in the absence and the presence of lysates from HeLa cells, visualized via the fluorescent label (6% native PAGE Cy3). 6% native PAGE WB shows immunostaining of proteins transferred to the PVDF membrane from the native PAGE gel, using α hnRNP K, α PCBP2, and α hnRNP A1 antibodies. (**C**) Loading control showing the initial amounts of proteins in the HeLa cell lysates, detected with SDS-PAGE and Western blotting.

#### Panel I

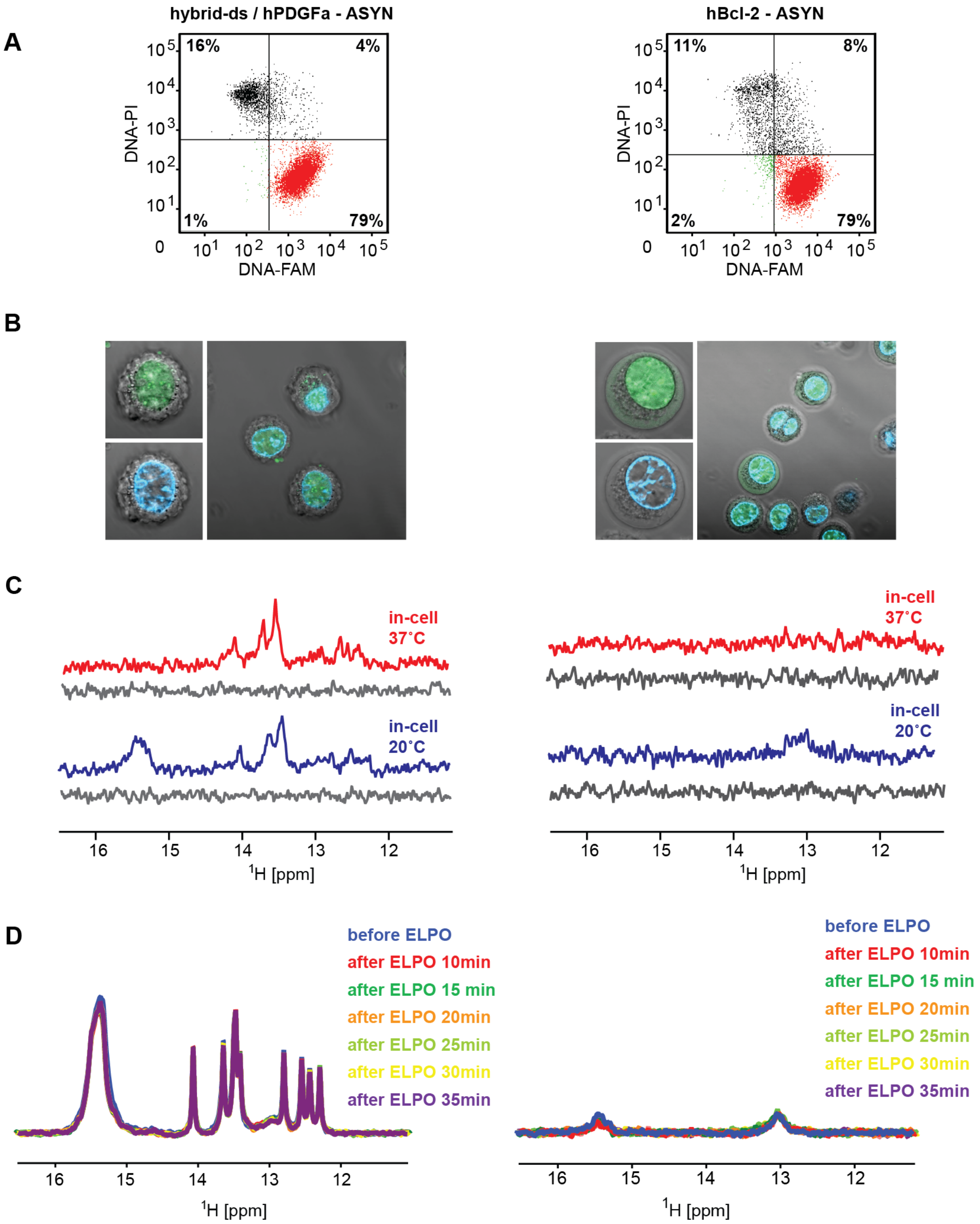

A
hybrid-ds / hPDGFa - ASYN
16%
4%
1%
79%
DNA-PI
DNA-FAM
hBcl-2 - ASYN
11%
8%
2%
79%
B
C
in-cell 37°C
in-cell 20°C
16 15 14 13 12
1H [ppm]
D
before ELPO
after ELPO 10min
after ELPO 15 min
after ELPO 20min
after ELPO 25min
after ELPO 30min
after ELPO 35min

**Panel II**

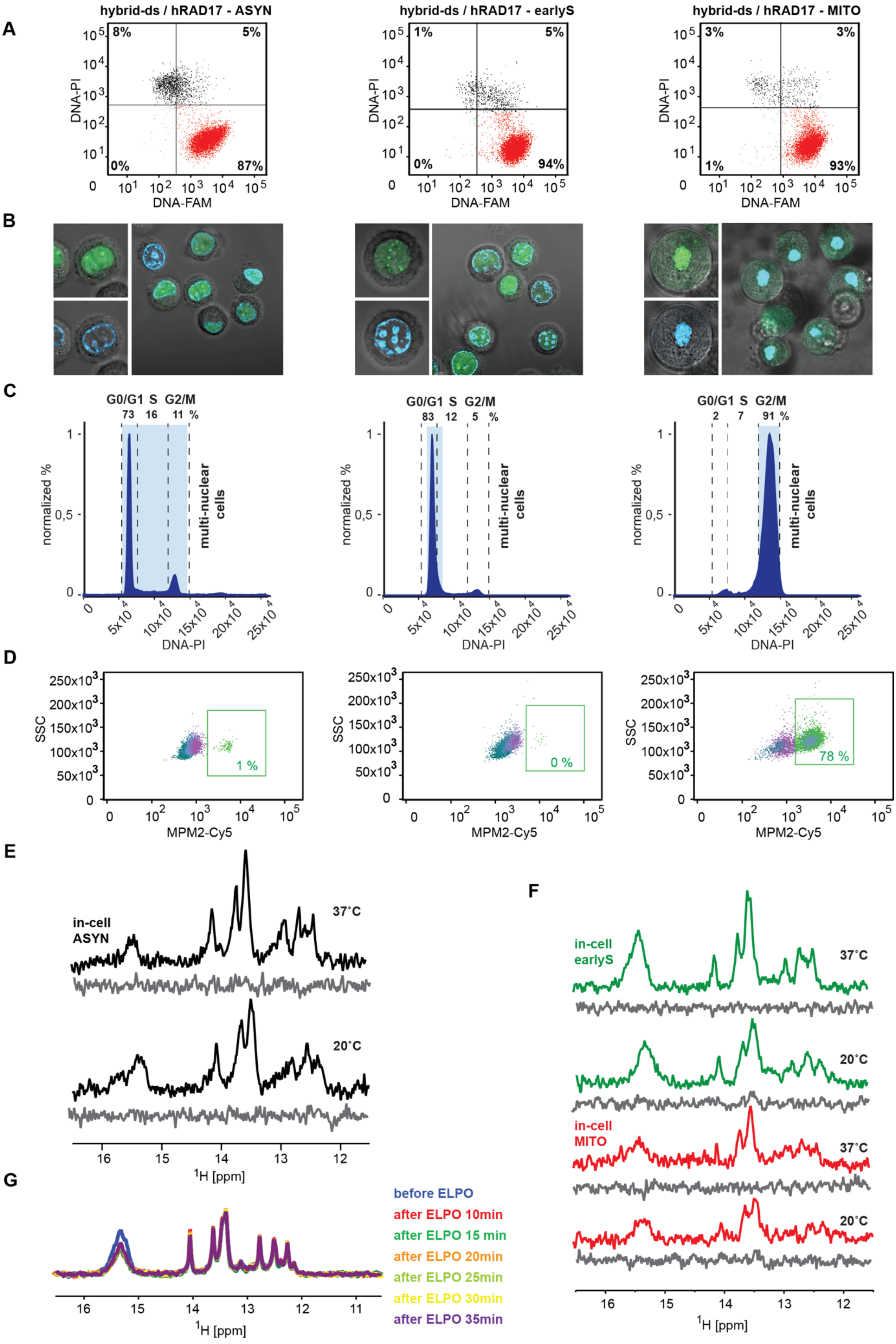

hybrid-ds / hRAD17 - ASYN
hybrid-ds / hRAD17 - earlyS
hybrid-ds / hRAD17 - MITO
A
B
C
D
E
F
G
DNA-PI
DNA-FAM
8% 5% 0% 87%
1% 5% 0% 94%
3% 3% 1% 93%
G0/G1 S G2/M
73 16 11 %
83 12 5 %
2 7 91 %
normalized %
multi-nuclear cells
SSC
MPM2-Cy5
1 %
0 %
78 %
in-cell ASYN
in-cell earlyS
in-cell MITO
37°C
20°C
$^1$H [ppm]
before ELPO
after ELPO 10min
after ELPO 15 min
after ELPO 20min
after ELPO 25min
after ELPO 30min
after ELPO 35min

**Fig. S7. Panel I - A**) Double-staining (DNA-PI/DNA-FAM) FCM analysis of asynchronous HeLa cells transfected with (FAM-)hybrid-ds/hPDGFa (left) and (FAM-)hBcl-2 (right). The FCM plots indicate the percentages of viable nontransfected cells, viable DNA-containing cells, dead/compromised nontransfected cells, and dead/compromised transfected cells in the bottom-left, bottom-right, top-left, and top-right quadrants, respectively. (**B**) shows confocal microscopy images of asynchronous HeLa cells transfected with (FAM-)hybrid-ds/hPDGFa (left) and (FAM-)hBcl-2 (right). Green and blue colors indicate the localization of FAM-labeled DNA and cell nucleus, respectively. (**C**) The imino regions of 1D $^1$H in-cell NMR spectra of hybrid-ds/hPDGFa (left) and hBcl-2 (right) acquired at 37 and 20 $^o$C in asynchronous cells. The corresponding regions of 1D $^1$H *in vitro* NMR spectra of the extracellular fluid taken from the samples after in-cell NMR spectra acquisition are shown in gray (leakage control). (**D**) shows an overlay of imino regions of 1D $^1$H *in vitro* NMR spectra of hybrid-ds/hPDGFa (left) and hBcl-2 (right) before and after applying the ”mock” electroporation pulse. **Panel II -** (**A**) Double-staining (DNA-PI/DNA-FAM) FCM analysis and (**B**) confocal microscopy images of asynchronous (left), early-S (middle) and M-synchronized (right) HeLa cells transfected with (FAM-)hybrid-ds/hRAD17. Green and blue colors indicate the localization of FAM-labeled DNA and cell nucleus/condensed chromatin, respectively. Synchronization controls: (**C**) Propidium iodide (PI) stained DNA content and (**D**) MPM2-Cy5 stained asynchronous (left), early-S (middle) and M-synchronized (right) HeLa cells transfected with (FAM-)hybrid-ds/hRAD17. The number in (**D**) indicates the percentages of mitotic cells in the population. (**E**) The imino regions of 1D $^1$H in-cell NMR spectra of hybrid-ds/hRAD17 acquired at 37 and 20 $^o$C in asynchronous cells and (**F**) early-S and M-synchronized Hela cells. The corresponding regions of 1D $^1$H *in vitro* NMR spectra of the extracellular fluid taken from the samples after in-cell NMR spectra acquisition are shown in gray (leakage control). (**G**) shows an overlay of imino regions of 1D $^1$H *in vitro* NMR spectra of hybrid-ds/hRAD17 before and after applying the ”mock” electroporation pulse.

**Panel I**

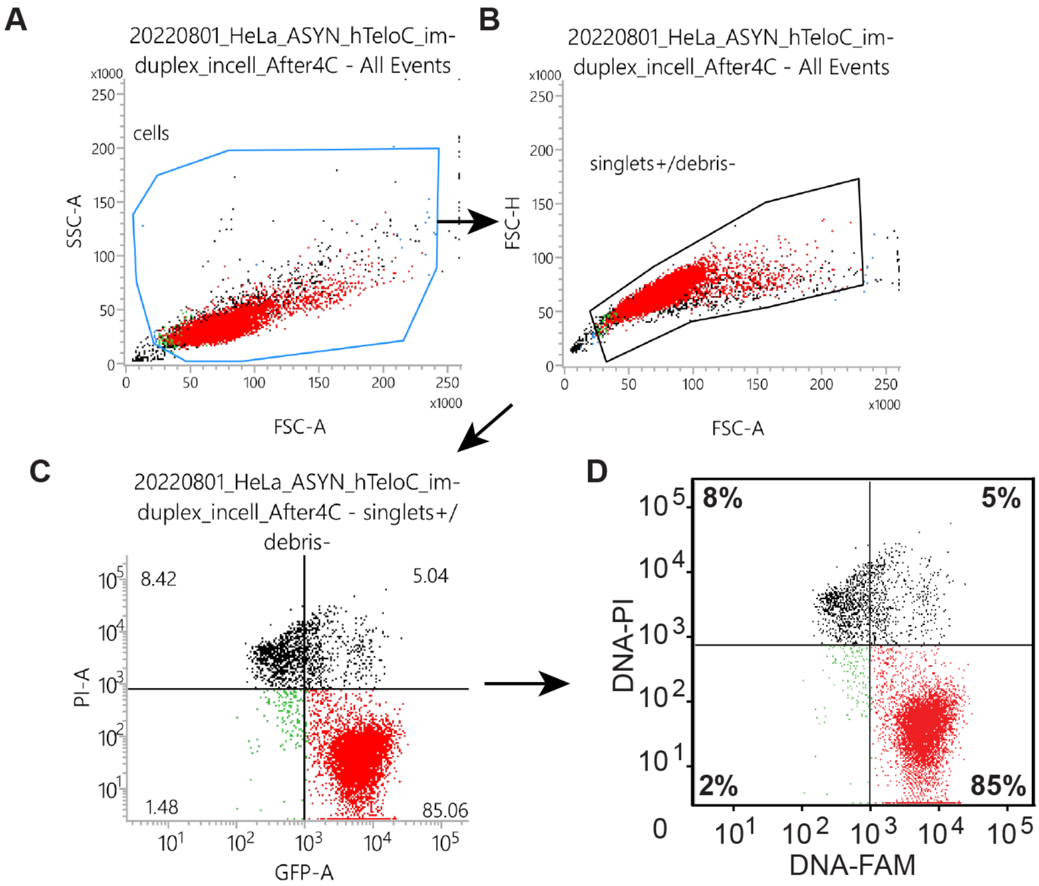
A
20220801_HeLa_ASYN_hTeloC_im-duplex_incell_After4C - All Events
cells
SSC-A
FSC-A
B
20220801_HeLa_ASYN_hTeloC_im-duplex_incell_After4C - All Events
singlets+/debris-
FSC-H
FSC-A
C
20220801_HeLa_ASYN_hTeloC_im-duplex_incell_After4C - singlets+/debris-
8.42
5.04
1.48
85.06
PI-A
GFP-A
D
8%
5%
2%
85%
DNA-PI
DNA-FAM

**Panel II**

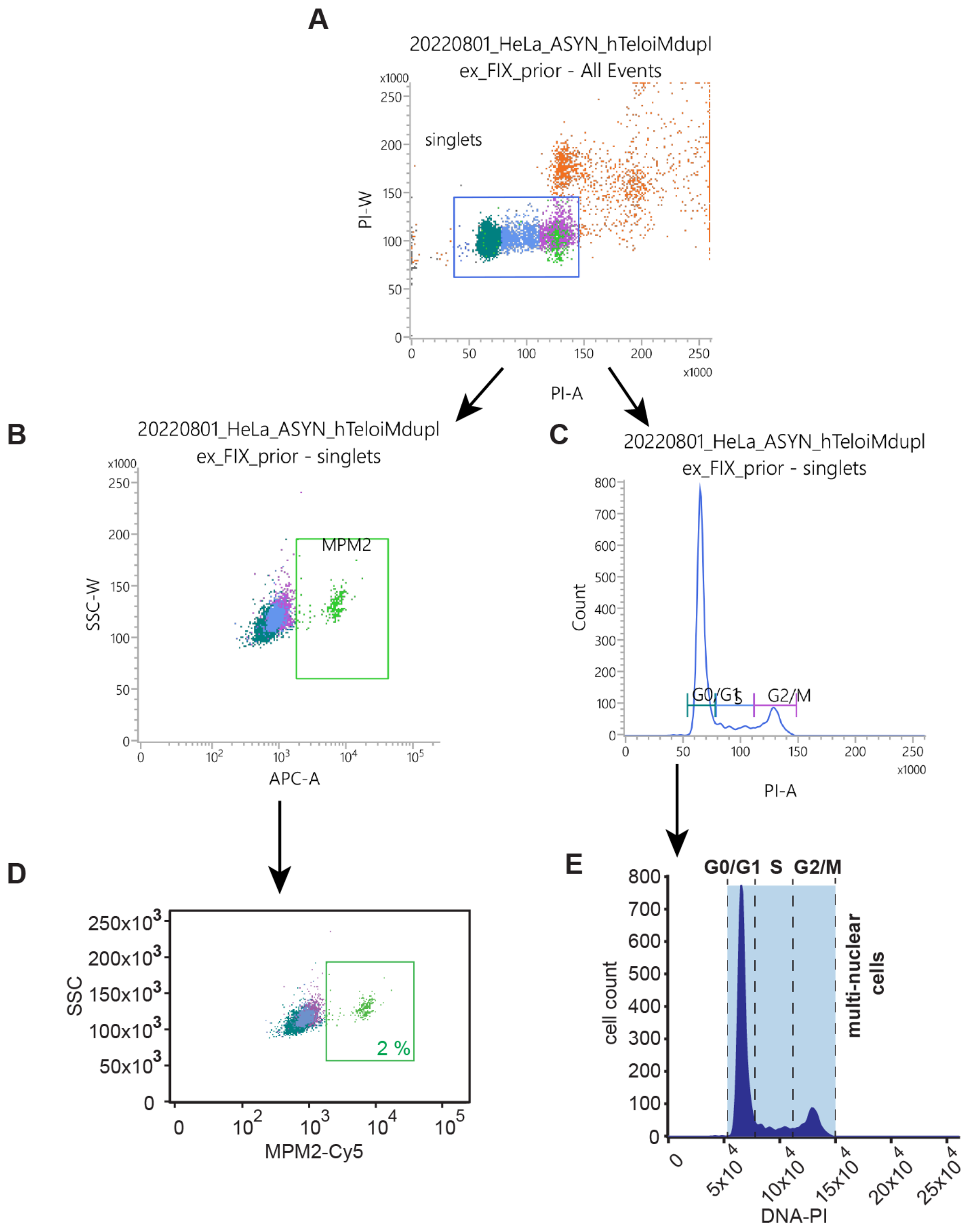


**Fig. S8. Panel I:** Example of FCS gating strategy to distinguish the apoptotic, dead cells or cells with compromised membrane integrity demonstrated on the FAM-hybrid-ds/hTel oligonucleotide transfected into asynchronous cells. (**A**) FSC-A and SSC-A gated cells to exclude the debris, (**B**) FSC-A/FSC-H gating to select singlet cells, (**C**) the final gating to visualize the cell viability (DNA-PI; y-axis) and the cell transfection efficiency (DNA-FAM; x-axis), and (**D**) the final processing by Adobe Illustrator CS6 V16.0.0. **Panel II:** Example of FCS gating strategy to evaluate cell cycle synchronization demonstrated on the hybrid-ds/hTel oligonucleotide transfected into asynchronous cells. (**A**) PI-A/PI-W gating to select singlet cells, (**B**) final gating to visualize the population of mitotic cells (MPM2-Cy5; x-axis), (**C**) final gating to distinguish individual phases of the cell cycle (DNA-PI; x-axis). (**D**) and (**E**) show the final processing by Adobe Illustrator CS6 V16.0.0 of (**B**) and (**C**), respectively.

**Table S2:** The potential technical issues impacting the in-cell NMR data interpretations regarding iM formation in-cells.

| | **Commentary** |
|---|---|
| *Perturbed metabolic homeostasis of the cells resulting from the overrepresentation of model iMFPS.* | Overrepresenting the DNA fragment in the intracellular space might alter the native cellular environment, particularly the molecular crowding (MC) levels in the intracellular space: MC is known to promote iM formation under *in-vitro* conditions. Admittedly, the in-cell NMR readout can be biased towards iM formation. Nonetheless, the fact that in-cell NMR observed iM equilibria resemble those observed in diluted solutions at neutral pH *in-vitro* suggests that potential bias stemming from artificially increased MC is small. Note: NMR detected (freely tumbling) exogenous DNA fragments' intracellular concentrations are in the range 10-20 μM. Total (freely tumbling + bound) exogenous DNA fragments' intracellular concentrations are estimated in the range 70-90 μM. |
| *Perturbed metabolic homeostasis of the cells resulting from the data acquisition in transfected pelleted cells.* | Pelleting cells might induce hypoxia connected with acidification of the intracellular space. To minimize the impact of the metabolic changes, which might occur on a time scale longer than 30 minutes,[38,39] we deliberately shortened the NMR acquisition window to about 10 min. Admittedly, our readout could still be biased towards iM formation. Nonetheless, the fact that we observed an equilibrium shifts in the opposite direction (iM unfolding) either suggests that hypoxia-induced artifacts are limited in the given acquisition time or that even hypoxia-induced acidification cannot override the main regulatory factor (the temperature) repressing iM formation in human cells. This explanation aligns with the outcome of the control bioreactor-based experiment performed for hybrid-ds/hT121-6 with a prolonged acquisition time of about 30 minutes (Fig S5). |
| *Restricted in-cell NMR data acquisition time.* | Some previously studied iMFPS displayed slow iM folding kinetics compared to their unfolding. A question arises whether the used acquisition time of 10 minutes upon temperature increase to 37 °C is enough for iMFPS to reach equilibrium. Admittedly, the readout could be biased towards iM unfolding. However, as demonstrated by the bioreactor-based in-cell NMR experiment for hybrid-ds/hT121-6, the respective iM remains unfolded in cells at 37 °C even during three times longer acquisition window, 30 minutes (cf. Fig. S5). Both acquisition times are well above the timescale of processes related to the postulated iM roles, transcriptional regulators, for instance. Therefore, to affect the binding of transcription factors to their consensus sequences or to stop/activate the transcription, an iM in the gene promoter should/would have to fold in a timescale close to the activity of the transcriptional machinery: Transcription factors bind their specific DNA sites on a time scale of seconds and their residence time varies from seconds to less than 10 minutes.[61,62] |

**Table S3**. The list of primary and secondary antibodies used in Shift-Westen-Blot assay.

*Primary antibodies*:

| Target | Origin | Dilution | Manufacturer | Cat. number |
|---|---|---|---|---|
| PCBP2 | rabbit polyclonal | 1:1000 | Origene | TA308051 |
| hnRNP K | mouse monoclonal | 1:1000 | Abcam | ab39975 |
| hnRNP A1 | mouse monoclonal | 1:1000 | Sigma Aldrich | R4528 |

*Secondary antibodies*:

| Target | Origin | Conjugated with | Dilution | Manufacturer | Cat. number |
|---|---|---|---|---|---|
| rabbit IgG | goat | HRP | 1:10000 | Jackson ImmunoResearch | 111-035-003 |
| mouse IgG | goat | HRP | 1:10000 | Jackson ImmunoResearch | 115-035-003 |
